# Cross-reactivity reduced dengue virus serotype 2 vaccine does not confer cross-protection against other serotypes of dengue viruses

**DOI:** 10.1101/480210

**Authors:** Jedhan Ucat Galula, Chung-Yu Yang, Brent S. Davis, Gwong-Jen J. Chang, Day-Yu Chao

## Abstract

The four serotypes of dengue virus (DENV) cause the most important rapidly emerging arthropod-borne disease globally. The humoral immune response to DENV infection is predominantly directed against the immunodominant cross-reactive weakly neutralizing epitopes located in the highly conserved fusion peptide of ectodomain II of envelope (E) protein (EDII_FP_). Antibodies recognizing EDII_FP_ have been shown to associate with immune enhancement in an *ex vivo* animal model. In this study, we explored how prime-boost strategies influence the immunogenicity of a cross-reactivity reduced (CRR) DENV-2 vaccine with substitutions in EDII_FP_ residues (DENV-2 RD) and found that mice in various DENV-2 RD prime-boost immunizations had significantly reduced levels of EDII_FP_ antibodies. In addition, heterologous DENV-2 RD DNA-VLP prime-boost immunization induced higher and broader levels of total IgG and neutralizing antibodies (NtAbs) although IgG titers to DENV-2 and 3 were statistically significant. Consistently, mice from DENV-2 RD DNA-VLP prime-boost immunization were fully protected from homologous DENV-2 lethal challenge and partially protected (60% survival rate) from heterologous lethal DENV-3 challenge. Our results conclude that the CRR DENV-2 RD vaccine requires a multivalent format to effectively elicit a balanced and protective immunity across all four DENV serotypes.

**Importance:** The low vaccine efficacy of the live-attenuated chimeric yellow fever virus-DENV tetravalent dengue vaccine (CYD-TDV) is unexpected and there is an urgent need to develop a next generation of dengue vaccine. Antibodies against the fusion peptide in envelope protein (E) ectodomain II (EDII_FP_) can potentially induce a severe disease via antibody-dependent enhancement (ADE) of infection. This study evaluated different formats of an EDII_FP_-modified DENV-2 vaccine (DENV-2 RD) in its capability of inducing a reduced EDII_FP_ antibodies, and sculpting the immune response towards an increased DENV complex-reactive neutralizing antibodies (CR NtAb). The results from this study confirmed the poor correlate of neutralizing assay with protection as suggested by the results of CYD-TDV clinical trials. There is a urgent need to develop a biological correlate with protection while evaluating the efficacy of the next generation dengue vaccine.

## Introduction

The four serotypes of dengue virus (DENV-1 to −4) are the most important causes of the rapidly emerging arthropod-borne disease in tropical and subtropical countries [1, 2], where an estimated 390 million dengue cases occur annually [3]. Infection with one serotype provides long-term homotypic protective immunity, while heterotypic protection is short-lived. Therefore, a tetravalent vaccine is necessary for a balanced immunity against all four DENV serotypes [4]. The most advanced dengue vaccine is the live-attenuated chimeric yellow fever virus-DENV tetravalent dengue vaccine (CYD-TDV) that expresses the premembrane (prM) and envelope (E) structural protein genes of each of the four DENV serotypes [5]. Even though pooled data from phase III clinical trials showed that CYD-TDV had lower efficacy against DENV-1 and −2 [6-8], it has been licensed in some countries under the condition of limited use for individuals aging 9 to 45 years old. A next-generation dengue vaccine with higher efficacy is urgently needed for use in a population of wider age group. Other candidates are in development including purified inactivated virus, subunit and DNA vaccines [9].

DNA vaccines afford advantages over the other vaccine platforms including ease and safety of production, elimination of replication interference [10], stimulation of CD8+ and CD4+ T cell responses similar to live-attenuated vaccines [11], and the possibility to formulate vaccines against multiple pathogens in a single vaccination. However, DNA vaccines elicit low neutralizing antibodies than protein-based vaccines due to unsatisfactory uptake by host cells and inadequate antigen expression. This was overcome by heterologous prime-boost immunizations, with combined use of DNA and other vaccine formats, which effectively induce a more robust and durable immune response against a number of medically important viruses including HIV [12], influenza [13, 14], HCV [15], and flaviviruses [16-22]. DENV infection elicits a poor quality of immune response that is highly skewed in the production of immunodominant antibodies against cross-reactive poorly neutralizing epitopes in E protein ectodomain II fusion peptide (EDII_FP_) [23, 24]. These EDII_FP_ antibodies can induce a severe disease resembling dengue hemorrhagic fever as demonstrated in AG129 mice via antibody-dependent enhancement (ADE) of infection [25-27]. Previously our group have constructed a DENV-2 wild-type (DENV-2 WT) plasmid containing the prM and E structural protein genes, which can be secreted outside of the transfected cells and forms virus-like particle (VLP) [28, 29]. The DENV-2 WT plasmid is also demonstrably immunogenic and protective in immunized mice when administered as DNA or VLP [28-30]. To circumvent the potential for a severe form of DENV-associated disease after infection by vaccine-induced EDII_FP_ antibodies through ADE, our group further engineered a cross-reactivity reduced (CRR) DENV-2 DNA vaccines with substitutions in the EDII_FP_ epitopes [24, 31]. These CRR DNA vaccines not only induced a significantly reduced levels of EDII_FP_ antibodies, which consequently reduced the enhancement of DENV infection *in vitro* and *in vivo*, but also sculpted the immune response towards an increased level of DENV complex-reactive neutralizing antibodies (CR NtAb) [32].

In this follow-up study, we evaluated if employing different prime-boost immunizations using the CRR EDII_FP_-modified DENV-2 vaccine (DENV-2 RD) can further broaden the CR NtAb responses and protect from infection by heterologous DENV serotypes.

## Materials and Methods

### Plasmid DNA and virus-like particle (VLP) production

Plasmids expressing DENV-2 strain 16681 prM and E structural protein genes, pVD2-WT (wild-type) and pVD2-RD (CRR, with substitutions in EDII_FP_ (G106R/L107D) were previously characterized in detail [24, 28, 29, 31]. Plasmids expressing DENV-1 and −3 (strains Panama), and −4 (strain Honduras) prM and E structural protein genes were previously characterized in detail [33]. Plasmids were produced in *E. coli* XL1-Blue cells (Stratagene, USA) and purified using Endofree Plasmid Maxi Kit (Qiagen, USA) according to the manufacturer’s instructions. VLPs were produced in COS-1 cells as previously described [34]. DENV-2 WT and RD VLPs were purified by 20% sucrose cushion and rate-zonal centrifugation on 5-25% sucrose gradients [35]. Protein concentration was determined by Bradford Assay (BioRad).

### Mice

Groups of 5 three-week-old ICR mice were immunized intramuscularly with 100 μg (50 μg in each thigh) of DENV-2 WT or RD DNA, or 4 μg of WT or RD VLP at 0 and 4 weeks, and bled retro-orbitally at 0, 4 and 12 weeks post-vaccination. The detailed grouping of mice for immunization is shown in **Supplemental Table 1**. The protective efficacy of DENV-2 RD vaccine was evaluated through passive transfer of maternal antibodies as previously described [28, 29]. Groups of two-day old ICR pups were obtained from immunized females mated with non-immunized males 3 weeks after boosting. Pups from females administered with TNE buffer served as challenge controls. DENV-2-specific NtAbs were confirmed prior to mating. Individual pups from each group were challenged intracranially with 10^6^ FFU equivalent to 141, 61, 11 and 1000-fold of 50% lethal doses (LD_50_) of DENV-1 Hawaii, DENV-2 16681, DENV-3 H87 and DENV-4 Honduras strains, respectively. Pups were monitored daily up to 21 days.

### ELISA

DENV-specific total IgG endpoint titers and EDII_FP_ epitope-specific IgG percentages were determined following the previously described Ag-capture ELISA [24, 29, 31]. All VLP antigens were standardized at a concentration producing an optical density at 450 nm (OD_450_) of 1.0. EDII_FP_ epitope-specific percentages were calculated as 100 × [1 – (RD VLP antigen endpoint/WT VLP antigen endpoint)]. The IgG1 and IgG2a isotype profiles of DENV-2-specific IgG in immune sera (diluted 1:500) were determined following the above Ag-capture ELISA protocol using the SBA Clonotyping System-HRP kit HRP-conjugated goat anti-mouse IgG1 and IgG2a (Southern Biotech, AL, USA). IgG avidities were also determined following the same Ag-capture ELISA with 6 M urea wash-step for 5 min at room temperature after incubation of pooled sera (diluted 1:100) with the antigen. The percent avidities were calculated as 100 × [(A_450_ VLP with urea) – A_450_ NC with urea)/(A_450_ VLP without urea – A_450_ NC without urea)] [36].

### Virus neutralization

Focus reduction micro-neutralization test (FRμNT) was performed as previously described [24, 29, 31]. The DENV strains used were DENV-1 Hawaii, DENV-2 16681, DENV-3 H87, and DENV-4 Honduras.

### Statistical analysis

One-way ANOVA followed by Tukey’s post-test was used for multiple comparisons. Data are means ± SD of two independent experiments.

## Results

### DENV-2 WT vaccine prime-boost immunizations elicit DENV-specific antibodies

To test if different prime-boost strategies using DNA or VLP would improve the immunogenicity of DENV-2 WT vaccine, ICR mice were immunized with combinations of DNA and VLP vaccines. All prime-boost strategies induced DENV-specific IgG titers ranging from 2.5 × 10^2^ to 3.7 × 10^5^ (**Fig. 1A-D**). The IgG mean titer against the homologous DENV-2 (**Fig. 1A, Supplemental Table 1)** was significantly higher with WT VLP-VLP (1.9 × 10^5^; p=0.0172) than WT DNA-DNA (2.3 × 10^4^). The titers induced by WT DNA-VLP or WT VLP-DNA were also slightly higher (8.3 × 10^4^ and 7.1 × 10^4^, respectively) than WT DNA-DNA without statistical significance. All prime-boost strategies induced IgG cross-reactive to heterologous DENV-1, −3 and −4. Again, WT DNA-DNA induced the lowest IgG titer against DENV-1 (**Fig. 1B**). Only WT VLP-DNA induced a significantly higher DENV-1 cross-reactive IgG mean titer (6.5 × 10^4^; p=0.0451) than WT DNA-DNA (1.3 × 10^4^), but not compared to WT VLP-VLP and WT DNA-VLP (1.8 × 10^4^ and 1.9 × 10^4^, respectively) (**Supplemental Table 1**). No significant differences were observed in DENV-3 (**Fig. 1C**) and DENV-4 (**Fig. 1D**) cross-reactive IgG titers.

**Fig. 1.**
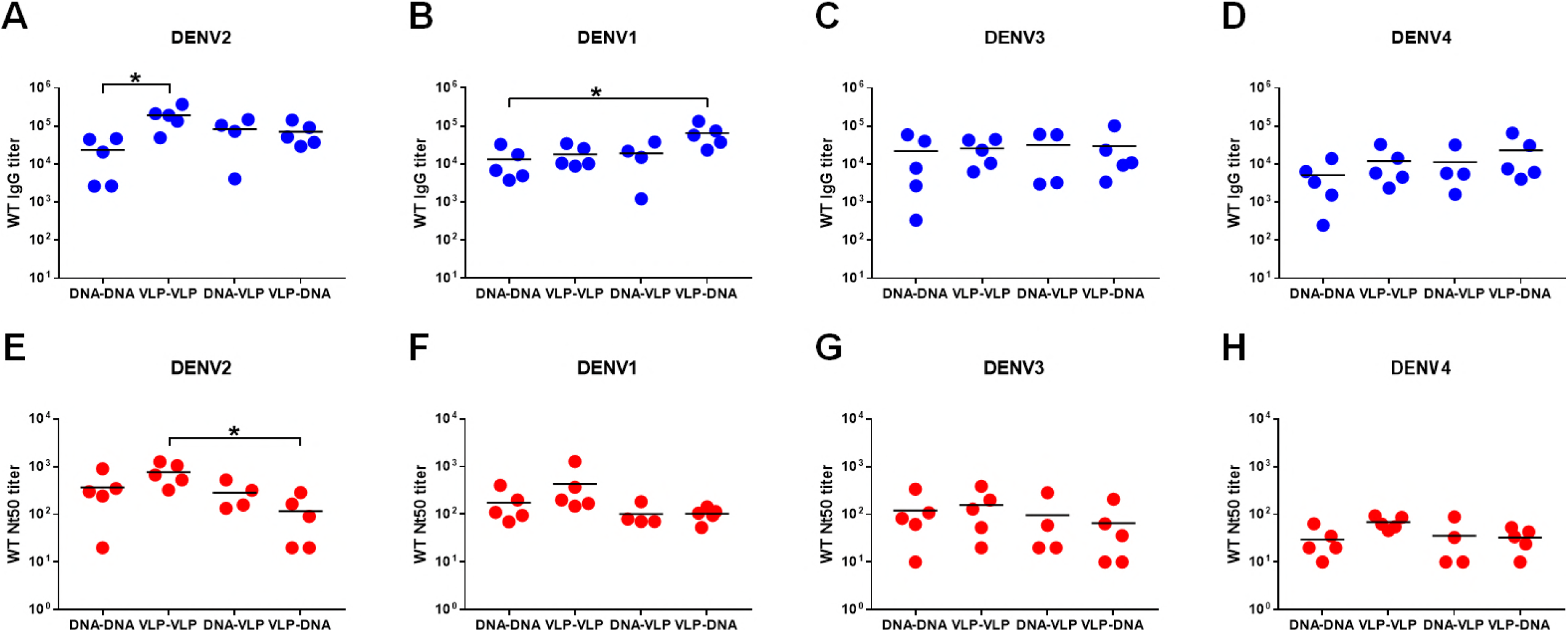
DENV-specific antibody responses induced by DENV-2 WT vaccine using various prime-boost immunization strategies. ICR mice (n = 5) were immunized twice at 0 and 4 weeks with different combinations of the WT DNA (100 µg) or VLP (4 µg) antigens. Immunization strategies separated by a dash represent priming with the first and boosting with the second (i.e., DNA-DNA, VLP-VLP, DNA-VLP; and VLP-DNA). Total IgG (**A-D**) and Nt_50_ (**E-F**) endpoint titers against homologous (**A, E**) DENV-2 and heterologous (**B, F**) DENV-1, (**C, G**) DENV-3, and (**D, H**) DENV-4 were determined by Ag-capture ELISA and 50% FRµNT. Multiple comparisons between groups were carried out on log-transformed data using one-way ANOVA with Tukey’s post-test. Horizontal bars indicate mean values. Data are means ± SD of two independent experiments. *, p<0.05.

All prime-boost strategies induced DENV-specific NtAb titers ranging from 10 to 1.3 × 10^3^ (**Fig. 1E-H**). Consistent with the total IgG results, WT VLP-VLP induced the highest NtAb titer to homologous DENV-2 (7.7 × 10^2^; **Fig. 1E, Supplemental Table 1**), although only significantly higher than WT VLP-DNA (1.2 × 10^2^; p=0.0169). Likewise, all prime-boost strategies induced antibodies with comparable cross-neutralizing activities against the heterologous DENV-1, −3 and −4 (**Fig. 1F-H**). These results demonstrate that heterologous WT DNA-VLP immunization induced a similar antibody response both in magnitude and neutralizing activity as homologous WT DNA-DNA and VLP-VLP immunizations against the homologous DENV-2 and the other heterologous DENV serotypes.

### CRR DENV-2 RD vaccine prime-boost immunizations elicit serotype-dependent CR NtAbs

We previously demonstrated that introducing targeted amino acid substitutions in EDII_FP_ of DENV-2 RD VLP had significantly ablated the reactivity of flavivirus group-reactive monoclonal antibodies such as 4G2 and 6B6C-1 [24], and immunization in mice with DENV-2 RD DNA circumvented the production of immunodominant weakly neutralizing and enhancing antibodies recognizing WT EDII_FP_ epitopes [31]. We then evaluated if DENV-2 RD antigens also dampen the immunodominance of the EDII_FP_ in the context of prime-boost immunizations. Consistent to the previous studies [24, 31], RD vaccines induced a significantly reduced proportions (means of 1%-6%) of IgG targeting the WT EDII_FP_ epitopes than the larger proportions (means of 32%-52%) induced by WT vaccines (**Fig. 2, Supplemental Table 1**). Comparably, WT-RD and RD-WT vaccine combinations also induced reduced WT EDII_FP_ IgG (means of 1%-18% and 1-28%, respectively) (**Fig. 2, Supplemental Table 1**).

**Fig. 2.**
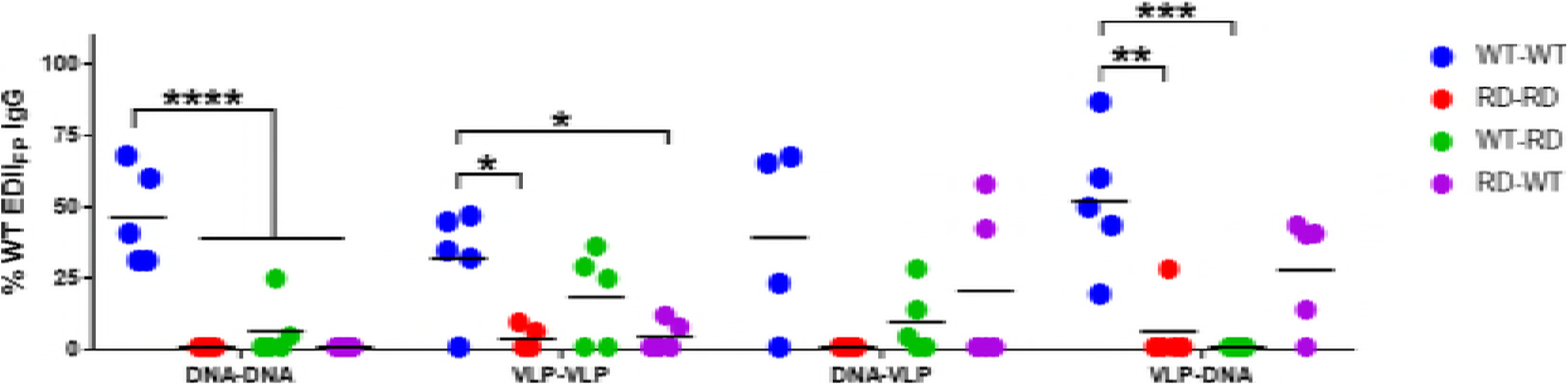
Percent WT EDII_FP_ epitope-specific IgG elicited by various prime-boost immunization strategies using different combinations of DNA and VLP antigens of the DENV-2 WT and RD vaccines. Multiple comparisons between groups were carried out using one-way ANOVA with Tukey’s post-test. Horizontal bars indicate mean values. Data are means ± SD of two independent experiments. *, p<0.05; **p<0.01; ***p<0.001; ****p<0.0001.

Next, we examined if different prime-boost strategies using DENV-2 RD vaccine can sculpt the immune response towards CR NtAbs as previously described [32] by comparing the immunogenicity against homologous and heterologous antigens. RD DNA-VLP induced consistently higher DENV-specific IgG titers to all four DENV serotypes (**Fig. 3A-D**). The IgG mean titer against the homologous DENV-2 (**Fig. 3A, Supplemental Table 1**) was significantly higher with RD DNA-VLP (4.9 × 10^5^) than RD VLP-VLP (3.7 × 10^4^; p=0.0072) and RD VLP-DNA (6.0 × 10^4^; p=0.0201), but not compared to RD DNA-DNA (6.6 × 10^4^). Consistently, the NtAb mean titer against the homologous DENV-2 was significantly higher with RD DNA-VLP (5.2 × 10^2^; **Fig. 3E, Supplemental Table 1**) than RD VLP-DNA (7.3 × 10^1^; p=0.0171) and RD DNA-DNA (3.2 × 10^1^; p=0.0125), but not compared to RD VLP-VLP (3.2 × 10^2^).

**Fig. 3.**
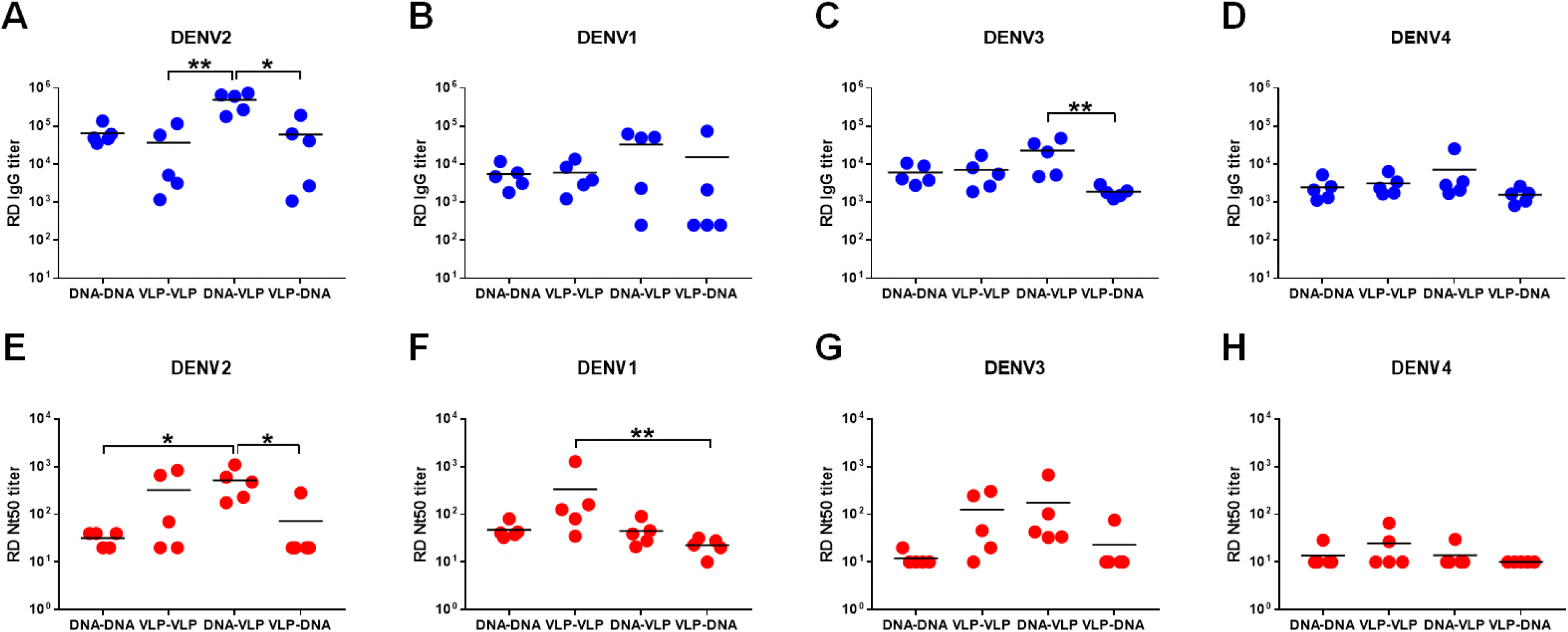
DENV-specific antibody responses induced by CRR DENV-2 RD vaccine using various prime-boost immunization strategies. ICR mice (n = 5) were immunized twice at 0 and 4 weeks with different combinations of the RD DNA (100 µg) or VLP (4 µg) vaccines. Immunization strategies separated by a dash represent priming with the first and boosting with the second (i.e., DNA-DNA, VLP-VLP, DNA-VLP; and VLP-DNA). Total IgG (**A-D**) and Nt_50_ (**E-F**) endpoint titers against homologous (**A, E**) DENV-2 and heterologous (**B, F**) DENV-1, (**C, G**) DENV-3, and (**D, H**) DENV-4 were determined by Ag-capture ELISA and 50% FRµNT. Multiple comparisons between groups were carried out on log-transformed data using one-way ANOVA with Tukey’s post-test. Horizontal bars indicate mean values. Data are means ± SD of two independent experiments. *, p<0.05; **p<0.01.

We further examined if prime-boost strategies using different forms of DENV-2 RD antigens would broaden the immune response to heterologous DENV serotypes by redirecting the immune response from the immunodominant EDII_FP_ site to DENV complex epitopes. Comparable DENV-1 and −4 (**Fig. 3B and D**) cross-reactive IgG titers were also induced, but a significantly higher DENV-3 cross-reactive IgG mean titer induced by RD DNA-VLP (2.3 × 10^4^; p=0.0023; **Fig. 3C, Supplemental Table 1**) than RD VLP-DNA (1.9 × 10^3^) was observed. On the contrary, the CR NtAb responses are serotype-dependent. A significantly higher DENV-1 CR NtAb mean titer was elicited by RD VLP-VLP (3.4 × 10^2^; p=0.0053; **Fig. 3F, Supplemental Table 1**) than RD VLP-DNA (2.3 × 10^1^). RD DNA-VLP elicited comparably higher CR NtAb mean titer against DENV-3 (1.8 × 10^2^; **Fig. 3G, Supplemental Table 1**). The CR NtAb mean titers to DENV-4 are equivalently low for all prime-boost strategies (<3.0 × 10^1^; **Fig. 3H, Supplemental Table 1**).

### CRR DENV-2 RD vaccine only protects against homologous DENV-2 infection

Next, we assessed if the CR NtAbs induced by different prime-boost immunizations of RD antigens can provide protection against lethal challenge of heterologous DENV serotypes. The lack of an ideal animal model to evaluate protective efficacy of a vaccine against the four DENV serotypes is one of the major hindrance in dengue vaccine development. Even though interferon α/β/γ receptor-deficient (AG129) mouse has been used in several studies for flavivirus vaccine evaluation [37-41], the challenge viruses for four serotypes of DENV were not available at the time when we started this study. We instead evaluated the efficacy of DENV-2 RD vaccine in lethally challenged suckling mice through passive protection by transferred maternal antibodies from immunized female mice as previously developed [28-30]. Utilizing suckling mouse model for evaluating vaccine efficacy allows us to evaluate the protection from dengue virus challenge only comes from maternal antibodies generated by vaccinated female mice and passively transferred to their babies. Pups from RD DNA-VLP, RD DNA-DNA, RD VLP-VLP, and RD VLP-DNA immunized mothers had 100%, 94%, 88%, and 83% protection, respectively, from homologous DENV-2 infection (**Fig. 4A**). However, no protection was observed after heterologous DENV-1, −3 and −4 infections (**Fig. 4B-D**), except for the pups from RD DNA-VLP immunized mother having 60% partial protection from DENV-3 challenge.

**Fig. 4.**
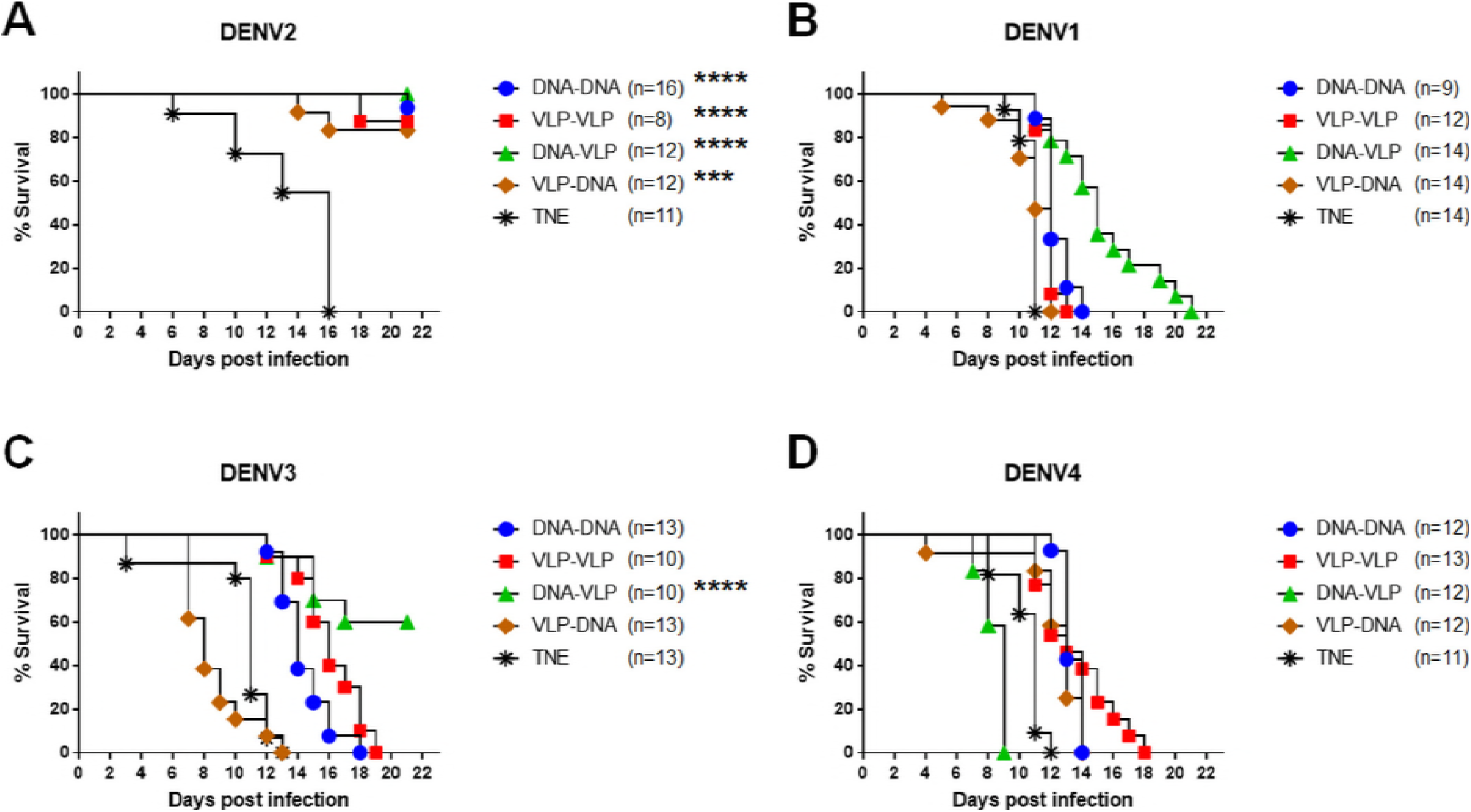
Protective efficacy of CRR DENV-2 RD vaccine using various prime-boost immunization strategies with different combinations of DNA and VLP antigens. Groups of two-day old pups from ICR female mice (Nt_50_ titer, >1:200) vaccinated with the designated DNA-DNA, VLP-VLP, DNA-VLP, and VLP-DNA prime-boost immunization strategies or TNE control were challenged intracranially with 10^6^ FFU of the homologous (**A**) DENV-2 (141-fold LD_50_) and heterologous (**B**) DENV-1 (61-fold LD_50_), (**C**) DENV-3 (11-fold LD_50_), and (**D**) DENV-4 (1000-fold LD_50_). Survival of the mice was monitored for 21 days after infection. The number (n) of pups in each group is indicated. Kaplan-Meier survival curves were analyzed by the log-rank test. (**E**) ***, p<0.001; ****, p<0.0001.

### CRR DENV-2 RD prime-boost immunizations elicit antibodies with varying isotypes and avidities

To determine if the absence of protection to heterologous DENV serotypes despite the induction of CR NtAb response by DENV-2 RD immunizations could be due to the functional quality of antibodies, we further measured the isotypes and avidities of IgG in sera from mice used in the immunogenicity experiment. RD DNA-DNA, DNA-VLP, and VLP-DNA induced largely IgG2a antibodies indicating a predominantly Th1 response, whereas RD VLP-VLP induced largely IgG1 antibodies indicating a predominantly Th2 response (**Fig. 5**). In addition, the elicited sera showed varying IgG avidities (**Supplemental Table 2**). RD DNA-VLP induced IgG with the highest avidity to the homologous DENV-2 followed by VLP-VLP, DNA-DNA, and VLP-DNA (51%, 50%, 33%, and 18% respectively). No avidities to DENV-1 and −3 were observed. Avidity to DENV-4 were also absent in sera elicited by RD VLP-VLP and VLP-DNA. However, RD DNA-DNA and DNA-VLP sera had minimal avidity to DENV-4 (21% and 3%, respectively) (**Supplemental Table 1**) yet had a very poor CR NtAb mean titers (<2.0 × 10^1^; **Fig. 3H**).

**Fig. 5.**
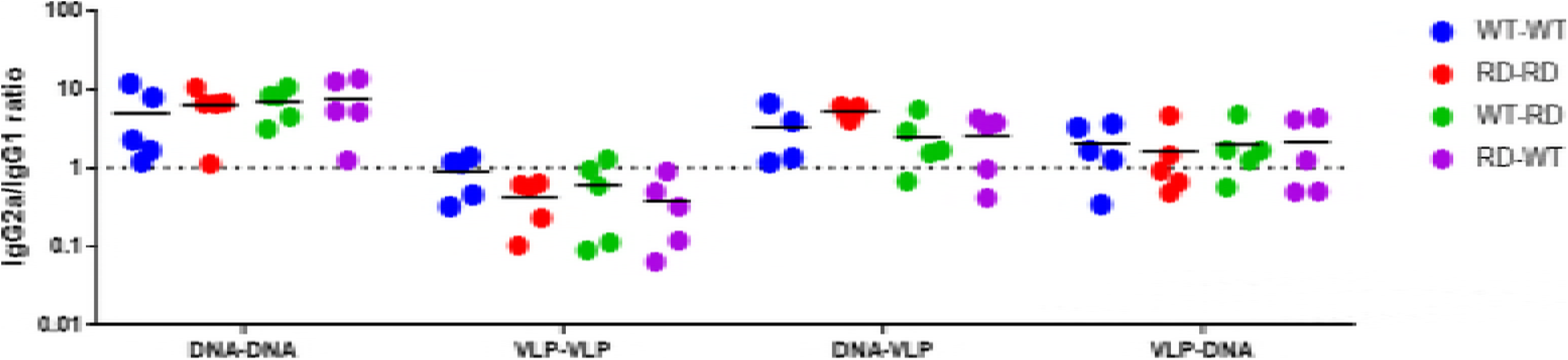
Comparison of IgG epitope profiles in mice immunized by various prime-boost immunization strategies using different combinations of DNA and VLP antigens of the DENV-2 WT and RD vaccines. The IgG isotype profiles were expressed as IgG2a/IgG1 ratios as indicator of Th1 (IgG2a/IgG1 ratio >1) and Th2 (IgG2a/IgG1 ratio <1) immune responses. Horizontal bars indicate mean values. Data are means ± SD of two independent experiments.

## Discussion

In this study, we extended our approach of dampening the immunodominance of potentially pathogenic EDII_FP_ epitopes by evaluating whether various prime-boost strategies using the DENV-2 RD vaccine can further broaden the CR NtAb response. Results demonstrate that heterologous RD DNA-VLP immunization can induce significantly higher levels of NtAbs to homologous DENV-2 serotype comparable to the homologous RD VLP-VLP immunization. This could be attributed to the efficient boosting capability of protein [19, 42-44], which is readily available for antigen presentation during immunization as opposed to the low-level expression of the immunogenic protein in response to DNA immunization. Moreover, RD VLP-VLP and DNA-VLP immunizations were able to induce an increase in DENV-1 and/or −3 CR NtAbs.

NtAbs has been considered to play a critical role in protection against DENV infection. Therefore, we also evaluated the antibody-mediated cross-protection elicited by DENV-2 RD vaccine through passive transfer of maternal antibodies. Interestingly, we detected CR NtAbs *in vitro* but it did not predict *in vivo* cross-protection. We observed that DENV-2 RD vaccine regardless of the prime-boost strategies is highly protective against homologous DENV-2 infection; however, it does not confer cross-protection against the heterologous serotypes. The elicited CR NtAbs evidently did not provide cross-neutralizing protection against the other DENV serotypes, which could be partly due to the amount of the elicited CR NtAbs and/or IgG avidities [45, 46]. As reports suggested, high antibody avidities correlate to better protection after vaccination [47, 48]. Indeed, the sera elicited from DENV-2 RD immunizations have avidities and high neutralization towards the homologous DENV-2 but not to the heterologous DENV-1 and −3. The absence of cross-protection against DENV-4 could be due to the suboptimal levels of DENV-4 CR NtAbs and/or the absence of avidities. In summary, our study confirms the poor correlate of neutralization titers with the protection from four serotypes of DENV infection as suggested by the results of CYD-TDV clinical trials [49, 50].

Overall, this study underscores the need for a multivalent formulation of our CRR DENV vaccines to fully harness its potential benefits with reduced ADE risk for dengue vaccine safety and broaden its protective efficacy. Importantly, the strategy of a heterologous DNA priming and VLP boosting highlights an effective platform of eliciting a better quality of immune response to DENV.

## Ethics Statement

This study complied with the guidelines for care and use of laboratory animals of the National Laboratory Animal Center, Taiwan. All animal experiments were approved by the Institutional Animal Care and Use Committee (IACUC) at the US Centers for Disease Control and Prevention (CDC), Division of Vector-borne Diseases (DVBD) and National Chung Hsing University (Approval Number: 101-58). All efforts were made to minimize suffering of mice.

## Author Contributions

JUG performed the experiments, data analyses, and wrote the paper. CYY performed the experiments and data analyses. BSD performed the experiments. GJC and DYC conceived the experiments, analyzed the data, and reviewed and edited the paper.

## Acknowledgments

This study was funded by grants 101-2321-B-005-018-MY2, 103-2321-B-005-014, and 104-2321-B-005-006- from the Ministry of Science and Technology, Taiwan.

